# The cyclin Cln1 controls polyploid titan cell formation following a stress-induced G2 arrest in *Cryptococcus*

**DOI:** 10.1101/2021.08.24.457603

**Authors:** Sophie Altamirano, Zhongming Li, Man Shun Fu, Minna Ding, Sophie R. Fulton, J. Marina Yoder, Vy Tran, Kirsten Nielsen

**Affiliations:** Department of Microbiology and Immunology, University of Minnesota, Minneapolis, Minnesota, USA; Cellular Imaging and Analysis Division, ThermoFisher Scientific, Coon Rapids, Minnesota, USA; Division of Pediatric Infectious Diseases, School of Medicine, Johns Hopkins University, Baltimore, Maryland, USA; Institute of Immunology & Immunotherapy, Institute of Microbiology & Infection, University of Birmingham, Birmingham, United Kingdom

## Abstract

The pathogenic yeast *Cryptococcus neoformans* produces polyploid titan cells in response to the host lung environment that are critical for host adaptation and subsequent disease. We analyzed the *in vivo* and *in vitro* cell cycles to identify key aspects of the *C. neoformans* cell cycle that are important for the formation of titan cells. We identified unbudded 2C cells, referred to as a G2 arrest, produced both *in vivo* and *in vitro* in response to various stresses. Deletion of the non-essential cyclin Cln1 resulted in over-production of titan cells *in vivo*, and transient morphology defects upon release from stationary phase *in vivo*. Using a copper-repressible promoter *P_CTR4_-CLN1* strain and a two-step *in vitro* titan cell formation assay, our *in vitro* studies revealed Cln1 functions after the G2 arrest. These studies highlight unique cell cycle alterations in *C. neoformans* that ultimately promote genomic diversity and virulence in this important fungal pathogen.

**Importance:** Dysregulation of the cell cycle underlies many human genetic diseases and cancers. Yet numerous organisms, including microbes, also manipulate the cell cycle to generate both morphologic and genetic diversity as a natural mechanism to enhance their chances for survival. The eukaryotic pathogen *Cryptococcus neoformans* generates morphologically distinct polyploid titan cells critical for host adaptation and subsequent disease. We analyzed the *C. neoformans in vivo* and *in vitro* cell cycles to identify changes required to generate the polyploid titan cells. *C. neoformans* paused cell cycle progression in response to various environmental stresses after DNA replication and before morphological changes associated with cell division, referred to as a G2 arrest. Release from this G2 arrest was coordinated by the cyclin Cln1. Reduced *CLN1* expression after the G2 arrest was associated with polyploid titan cell production. These results demonstrate a mechanism to generate genomic diversity in eukaryotic cells through manipulation of the cell cycle that has broad disease implications.

## Introduction

Cell division during mitosis typically involves a single round of DNA replication and then equal partitioning of this DNA into two separate, genetically identical cells. Despite the highly regulated process of cell division, cell cycle alterations that produce whole genome duplications occur naturally throughout the tree of life (1). These whole genome duplications produce polyploids, which are cells with increased DNA content beyond the typical ploidy state of the organism. Polyploidy is a normal part of development, tissue homeostasis, and stress response in many multicellular eukaryotes (2). For example, trophoblast giant cells found in mammalian placenta duplicate their genomes while bypassing mitosis and cytokinesis, a process known as endocycling, to form polyploid cells that can exceed 512C (3). Studies in *Drosophila* show that puncture wound healing of the epithelium involves cell fusion and endocycling, both of which produce polyploid cells (4). Additionally, keratinocytes are known to polyploidize, possibly increasing the potential for cellular survival during UV exposure while decreasing the oncogenic potential of the cells (5–8). Unfortunately, while evidence for polyploidy across many kingdoms of life has been identified, the biological function of polyploidy in many cases is still unknown.

Polyploid cells are often not stable and can undergo chromosomal loss or aberrant mitosis to produce cells with an abnormal number of chromosomes that are not an integer multiple of the base genome, known as aneuploid cells (9, 10). The ability to form polyploid and aneuploid cells can be beneficial to an organism but can also make an organism more susceptible to disease. Polyploidy is often associated with the formation and progression of cancer cells due to this ability to generate aneuploid cells. Approximately 90% of solid tumors and 75% of hematopoietic cancers are aneuploid (11) with recent models suggesting that these aneuploid cells likely arise from polyploid precursors (9, 12–14).

The molecular mechanisms underlying polyploid division and how polyploidy contributes to genomic instability still remain largely unknown. A better understanding of this process would provide insight into the pathological role of polyploidy. The study of polyploidy has generally been constrained to multicellular organisms and genetically engineered yeast. Polyploidy in multicellular organisms is a complex multifactorial process involving temporal and tissue specific cues that make replicating these processes in a laboratory setting challenging (8). Previous studies in yeast have been very helpful to define the genomic instability of polyploids but required genetically modified strains to induce and study polyploidy (15–17). Recently, the human pathogenic yeast *Cryptococcus neoformans* was found to produce enlarged, polyploid cells, known as titan cells, during pulmonary infections (18, 19).

*C. neoformans* is a haploid (1C) budding yeast with cells that are typically 5-7 µm in cell diameter. In contrast, the titan cells have ploidies ranging from 4C-312C and are 15-100 µm in cell diameter (18, 19). Titan cells undergo cell division to produce genetically distinct 1C or aneuploid daughter cell populations that are typical-sized and exhibit increased stress resistance, suggesting that the production of titan cells during infection promotes rapid adaptation to the host environment (20). In addition, titan cells have cell wall changes that impact the host immune response (21, 22). Due to the vast size difference between typical and titan cells, these *C. neoformans* cells can be readily separated and are a good model system to study the mechanism of naturally occurring polyploid cell formation and subsequent error-prone cell division that leads to genetic variation.

Here we characterize the *C. neoformans* cell cycle during infection and *in vitro* to identify how regulation of the cell cycle in *C. neoformans* leads to titan cell formation and the resulting polyploidy. Using homology to known cyclins and cyclin dependent kinases (CDKs), we identified the cyclin, Cln1, as a master regulator of titan cell formation. We show *C. neoformans* has a G2 arrest with no bud formation in response to the murine pulmonary environment and also under *in vitro* nutrient deprivation conditions. We show that Cln1 is critical for balancing DNA replication and cell division after the G2 arrest. When *CLN1* expression is low, polyploid titan cells are produced in response to specific environmental stimuli. Taken together, these studies identify Cln1 as a master regulator underlying ploidy changes in *C. neoformans* and highlight the versatility of this unicellular eukaryotic microorganism to understand the molecular regulation and biological function of ploidy changes.

## Results

### *C. neoformans* has an unbudded G2 arrest during *in vivo* infection and *in vitro* nutrient deprivation

Cell cycle progression is often correlated with cell size and shape (23), and thus cell cycle changes likely precede the observable size and morphology changes that produce the titan cells. The mitotic cell cycle is typically divided into four phases: growth 1 (G1) prior to DNA replication; synthesis (S) when the DNA is replicated; growth 2 (G2) prior to cell division; and mitosis (M) when cytokinesis occurs and the cell divides. *C. neoformans* is a budding yeast, similar to the model yeast *Saccharomyces cerevisiae*. Previous studies in *S. cerevisiae* linked distinct cell morphologies with the four cell cycle phases (23). In *S. cerevisiae*, G1 cells are unbudded and have a 1C DNA content. A bud appears at the beginning of S phase and grows as the DNA is replicated. Entry into G2 occurs when DNA synthesis is completed, and the daughter cell is approximately half the size of the mother cell. M phase involves cytoskeletal tubulin rearrangements to generate the mitotic spindle that pulls the replicated DNA into the daughter cell (24). Similar cell morphology and ploidy changes associated with cell cycle phases were previously shown to occur in log phase cultures of *C. neoformans in vitro* (25, 26) and are consistent with a cell cycle progression similar to that in *S. cerevisiae*.

Yet our previous study showed that *in vivo* typical sized cells were primarily 2C (18), instead of the predicted mixed population of 1C and 2C cells observed in log phase cultures grown *in vitro*. To define the *in vivo* cell cycle that precedes titan cell formation, we isolated typical sized cells based on DNA content by fluorescence-activated cell sorting (FACS), assessed the morphology of each sorted population, and analyzed the ploidy and morphology compared to that of log phase and stationary phase cells grown *in vitro* (**Figure 1A**). As shown previously (26), our *C. neoformans* log phase cells consisted of 1C and 2C peaks, where the majority of the 1C population was unbudded (∼96%, n=226) and the majority of the 2C population was budded (∼81%, n=230) (**Figure 1A**). Also consistent with previous studies (26, 27), the stationary phase cells generated via nutrient starvation arrested after DNA synthesis to produce a population that consisted primarily of 2C unbudded cells (∼98%, n=215) (**Figure 1A**). The *in vivo* typical sized cell population consisted of both 1C and 2C populations; the 1C population was ∼ 89% unbudded (n=357), while the 2C population was ∼93% unbudded (n=203) (**Figure 1A**). Similar to the stationary phase 2C cells, the *in vivo* typical sized 2C cell population was primarily unbudded.

**Figure 1.**
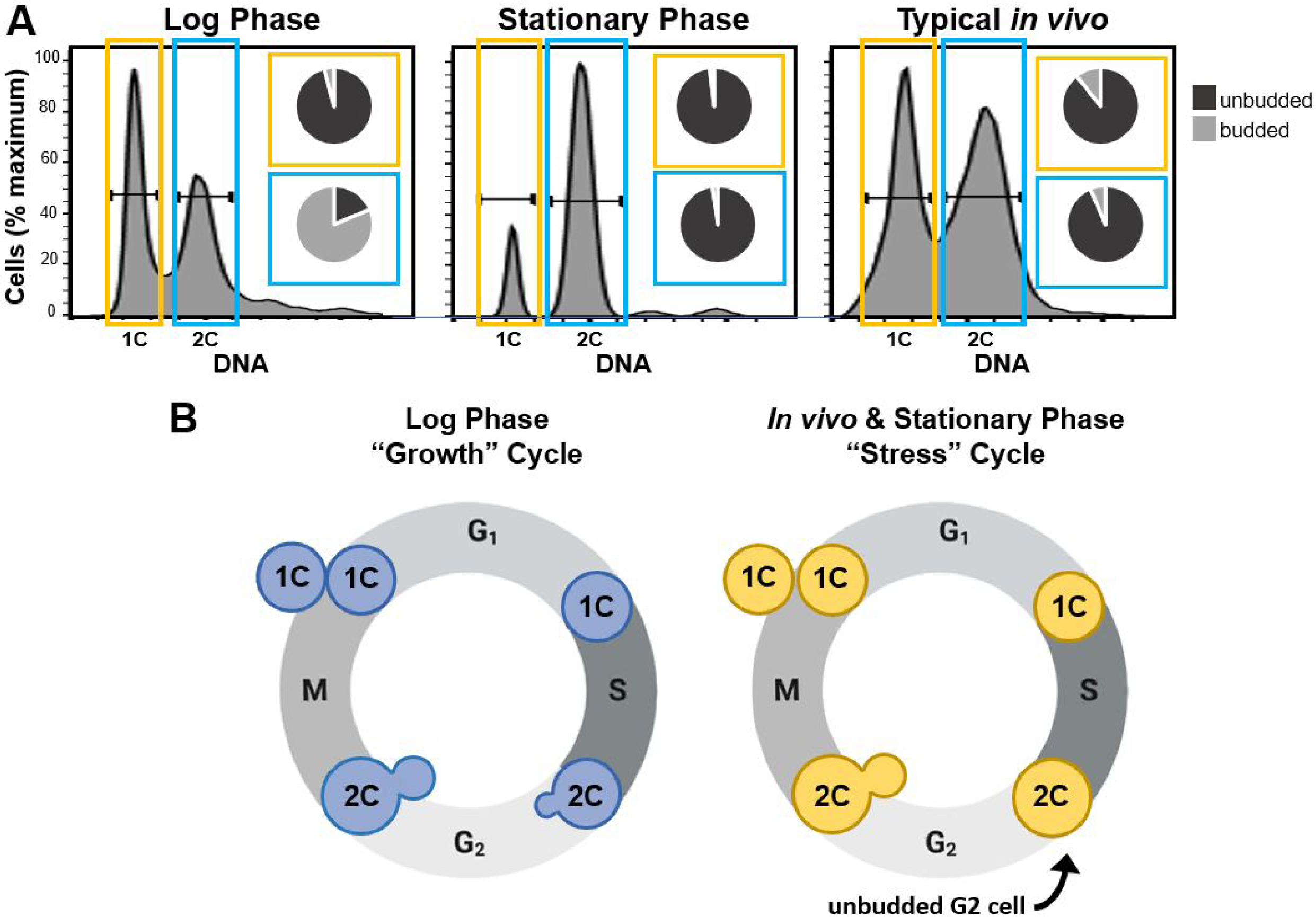
*In vivo* typical sized cells and *in vitro* stationary phase cells are primarily unbudded 2C cells. A) Analysis of cell morphology with 1C (yellow) and 2C (blue) DNA content in log phase cells, stationary phase cells, and typical size cells isolated from the lungs of mice at 14 days post infection (typical *in vivo*). Cells were grown in the indicated conditions, fixed, stained with propidium iodide, and then fluorescence activated cell sorting (FACS) was performed to purify the 1C and 2C cell populations. The resulting purified cells were then analyzed microscopically to determine the proportion of cells containing buds (insets). B) Schematic representation of the *C. neoformans* “growth” cell cycle (blue cells) observed in log phase cells and the putative “stress” cell cycle (yellow cells) that is notable for production of unbudded 2C cells that were observed in both stationary phase cultures *in vitro* and among typical sized cells *in vivo*.

Our results show the *C. neoformans* log phase cells grown in nutrient replete media produced a 2C population that was primarily budded, similar to the cell cycle observed in *S. cerevisiae*, where DNA replication is associated with daughter cell growth and division (**Figure 1B, “Growth” Cycle**). In contrast, the *C. neoformans* stationary phase cells and *in vivo* typical sized cells produced a 2C population consisting of unbudded cells. These data show *C. neoformans* cells in both *in vivo* and *in vitro* (nutrient, hypoxia, temperature) (26–29) “stress” conditions utilize an alternative cell cycle in which DNA replication occurs before any morphological changes associated with daughter cell growth, resulting in an unbudded G2 arrest (**Figure 1B, “Stress” Cycle**).

### Nuclear dynamics are predominantly normal upon cell cycle reentry after unbudded G2 arrest

We envisioned two possible scenarios in response to the unbudded G2 arrest. One hypothesis was that the unbudded G2 arrest is a cell cycle anomaly and results in an aberrant mitosis upon reentry into the cell cycle. Alternatively, we hypothesized that the unbudded G2 arrest is a pre-programed part of the *C. neoformans* cell cycle. In this scenario, we expected to see little to no disruption in mitosis, with the cells readily adapting to the altered “stress” cell cycle and 2C (or higher for titan cells) DNA content.

To test these two hypotheses, we analyzed nuclear dynamics and movement upon cell cycle reentry through analysis of cells expressing fluorescently-tagged versions of the proteins Nop1 (nucleolus), Tub1 (tubulin), Cse4 (inner kinetochore), and Ndc1 (nuclear envelope) (**Figure 2**). *In vitro* log phase, *in vitro* stationary phase, and *in vivo* titan cells were suspended in nutrient replete medium to induce re-entry into the cell cycle and analyzed by time-lapse microscopy. Surprisingly, in all cell types only very subtle differences from the log phase cells were observed. As shown previously, in the log phase cells the nuclear membrane ruptures, resulting in loss of nucleolus staining as the mitotic spindle elongates into the daughter cell, followed by half of the centromeres translocating back into the mother cell (25). (**Figure 2A**). The dynamics of this process were very similar in the *in vitro* stationary phase (**Figure 2B**) and *in vivo* titan cells (**Figure 2C**). The only observable difference was a delay in loss of nucleolus staining. This was most obvious in the stationary phase cells where the nucleolus staining diminished dramatically but did not fully disappear (**Figure 2B**). A similar phenomenon was observed during titan cell division, where nucleolus signal loss was slightly delayed compared to the log phase cells (**Figure 2C**). It should be noted that both the stationary phase and titan cells are larger than the log phase cells and have a higher ploidy, thus the delay in loss of nucleolus staining could be due to either a larger nucleus or cell that results in more time before rupture of the nuclear membrane by the spindle. Taken together, we observed no increased rates of spindle, chromosome localization, or morphological anomalies such as trimeras or elongated buds that would suggest an aberrant mitosis upon reentry into the cell cycle following the unbudded G2 arrest, suggesting the unbudded G2 arrest was a programmed cell cycle response to stress in *C. neoformans*.

**Figure 2.**
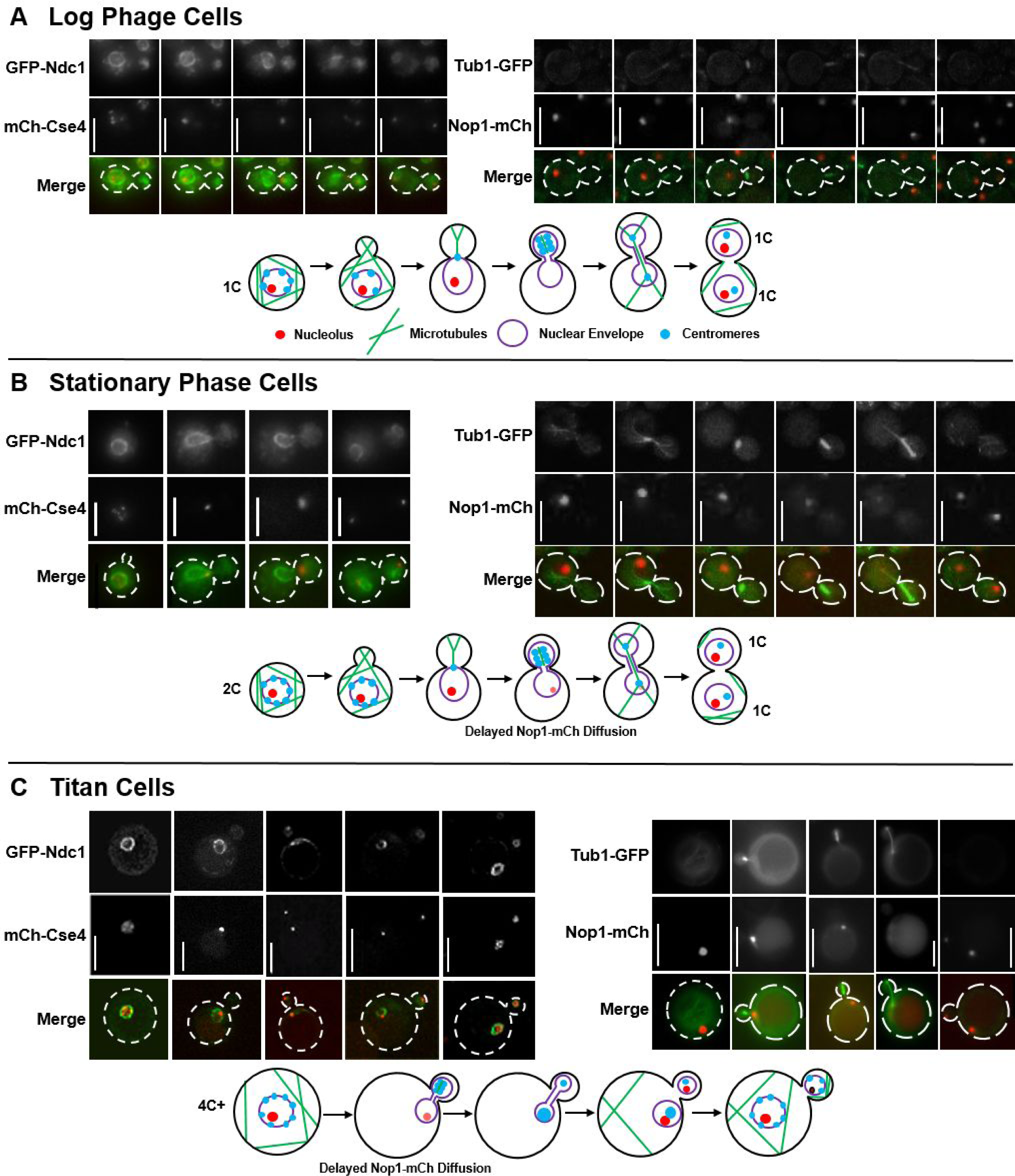
Analysis of nuclear dynamics after unbudded G2 arrest shows minimal effect on the ability of stationary and titan cells to complete mitosis and divide. Representative images and corresponding schematic diagrams showing mitotic events in log phase cells (A), stationary phase cells (B) and *in vivo* titan cells (C). Centromere and nuclear envelope dynamics were analyzed in cells expressing the inner kinetochore protein, mCherry-Cse4, and the nuclear envelope protein, GFP-Ndc1. Microtubules and nucleolus dynamics were analyzed in cells expressing the nucleolar protein, Nop1-mCherry, and the tubulin protein, Tub1-GFP. All cells were analyzed by time-lapse microscopy and titan cells were analyzed using both time-lapse and 2-photon microscopy due to their larger size. Not all representative images of the different cell cycle stages are of the same cell. A) Consistent with previous studies of log phase cells (25), the centromeres were not clustered in the mother cell prior to mitosis, microtubules formed the mitotic spindle in the daughter cell and the centromeres then clustered and moved completely into the daughter, elongation of the spindle resulted in breakage of the nuclear envelope and loss of the nucleolus staining. After mitosis, half of the centromeres returned to the mother cell, two separate nucleoli and two separate nuclear envelopes were formed – one in the mother and one in the daughter cell. B) Stationary phase cells that had already undergone DNA replication prior to bud emergence underwent mitosis similar to log phase cells. A minor difference in nucleolus retention time was observed. The nucleolus of the stationary phase cells faded dramatically, but was retained throughout the course of mitotic spindle formation and DNA segregation. C) *In vivo* derived titan cells also underwent a mitosis similar to the log and stationary phase cells, including a slightly delayed disappearance of the nucleolus in the mother cell. The titan cells produced typical sized daughter cells. Scale bars are 5 µm in panels A and B and 10 µm in panel C.

### The non-essential cyclin Cln1 negatively regulates *in vivo* titan cell formation

In other human pathogens DNA replication is linked to cell growth and division; and changes in cell cycle progression affect cell morphology (30–32). Thus, we hypothesized that the unbudded G2 arrest and subsequent polyploid titan cell formation is also regulated by the cell cycle.

To explore the role of cell cycle regulation in titan cell formation, we identified 14 putative cyclins and seven cyclin dependent kinases (CDKs) in the *C. neoformans* genome based on homology to *S. cerevisiae* and mammalian cyclins and CDKs (**Table 1**). Deletion strains were generated in the KN99α genetic background for each of the *C. neoformans* cyclin and CDK homologs and analyzed for their effect on the *in vitro* cell cycle via ploidy and morphology analysis (**Supplementary Figure SF1A and SF1B**). We were unable to obtain deletions for two cyclins (CLB2 and CCL1) and four CDKs (CDK1, SGV1, KIN28, and PHO85), suggesting that these cyclins and CDKs are essential in the KN99α strain. This essentiality was further supported by diploid sporulation assays (**Supplementary Table ST1**). The expression of *CLB2* and *CCL1* was used to determine when these cyclins likely functioned within the *in vitro C. neoformans* cell cycle (**Supplementary Figure SF1C)**.

**Table 1.** Identification of putative cyclins and cyclin dependent kinases in *C. neoformans*.

| Function<sup>†</sup> | Gene ID | Name | Essential<sup>‡</sup> | Cyclin Family |
| --- | --- | --- | --- | --- |
| Cyclins | CNAG_06092 | Cln1 | No | Cdc28 |
|  | **CNAG_04575** | **Clb2** | **Yes** | Cdc28 |
|  | CNAG_02095 | Clb3/Cbc1 (79) | No | Cdc28 |
|  | CNAG_00183 | Pcl2 | No | Pho85 |
|  | CNAG_00442 | Pcl9 | No | Pho85 |
|  | CNAG_03385 | Pcl103 | No | Pho85 |
|  | CNAG_02658 | Pcl5 | No | Pho85 |
|  | CNAG_05524 | Pcl7 | No | Pho85 |
|  | CNAG_01922 | Pho80 | No | Pho85 |
|  | CNAG_00024 | Clg1 | No | Pho85 |
|  | CNAG_00440 | Ssn801 | No | Ssn3 |
|  | CNAG_02127 | Ssn802 | No | Ssn3 |
|  | CNAG_05901 | Ssn803 | No | Ssn3 |
|  | **CNAG_04405** | **Ccl1** | **Yes** | Kin28 |
| Cyclin dependent kinases | **CNAG_01664** | **Cdk1** | **Yes** |  |
|  | CNAG_04118 | Ctk1 | No |  |
|  | **CNAG_05549** | **Sgv1** | **Yes** |  |
|  | **CNAG_06445** | **Kin28** | **Yes** |  |
|  | **CNAG_08022** | **Pho85** | **Yes** |  |
|  | CNAG_00415 | Cdc2801 | No |  |
|  | CNAG_06086 | Cdk8 | No |  |

<sup>†</sup>Function of predicted protein product<sup>‡</sup>Essential cyclins and CDKs are in bold and were determined based on diploid sporulation assays (see ST1 for further detail)

To assess titan cell formation in each of the deletion strains, we infected mice with the deletion strain or the corresponding complement strain and assessed the ability of each to produce titan cells (**Supplementary Figure SF2**). The *clb3Δ* and *ssn803Δ* mutants had severe 37°C growth defects and cells from mice were insufficiently recovered to analyze their titan cell formation (data not shown and **Supplementary Figure SF2**). Deletion of the cyclins Pcl5, Clg1, Ssn802, and the CDK Ctk1 had subtle effects on titan cell formation that were rescued by complementation.

Interestingly, deletion of the cyclin gene, *CLN1,* led to 98% ± 0.27% titan cell formation (**Figure 3A**, p=0.0000016), while the complement exhibited titan cell production similar to wild type. To further investigate the role of *C. neoformans* Cln1 in titan cell production, we generated two *CLN1* overexpression strains. One overexpression strain with *CLN1* under the control of the constitutive promoter, *GPD1* (33), and a second overexpression strain with *CLN1* under the control of the copper repressible promotor, *CTR4*. The *CTR4* promoter is turned on in the absence of copper and turned off in the presence of copper (34). The lung environment is a copper-limiting environment (35), resulting in overexpression of *CLN1* in the *P_CTR4_-CLN1* strain *in vivo*. *CLN1* overexpression reduced titan cell production compared to the wild type strain (**Figure 3A**, p=0.000011). The *cln1Δ* cells had a significant increase in cell body diameter compared to the wild type cells, while the overexpression strains had cell body sizes smaller than the wild type cells (**Figure 3B**). In addition to their large size, titan cells are defined by their increased ploidy and the high chitin content in their cell wall (36–38). The large *cln1Δ* cells isolated from mice had higher levels of both DNA and chitin, indicating the *cln1Δ* cells are bona fide titan cells (**Figure 3C and 3D**). Overall, these data show *CLN1* gene expression negatively regulates titan cell formation during *in vivo* infection.

**Figure 3.**
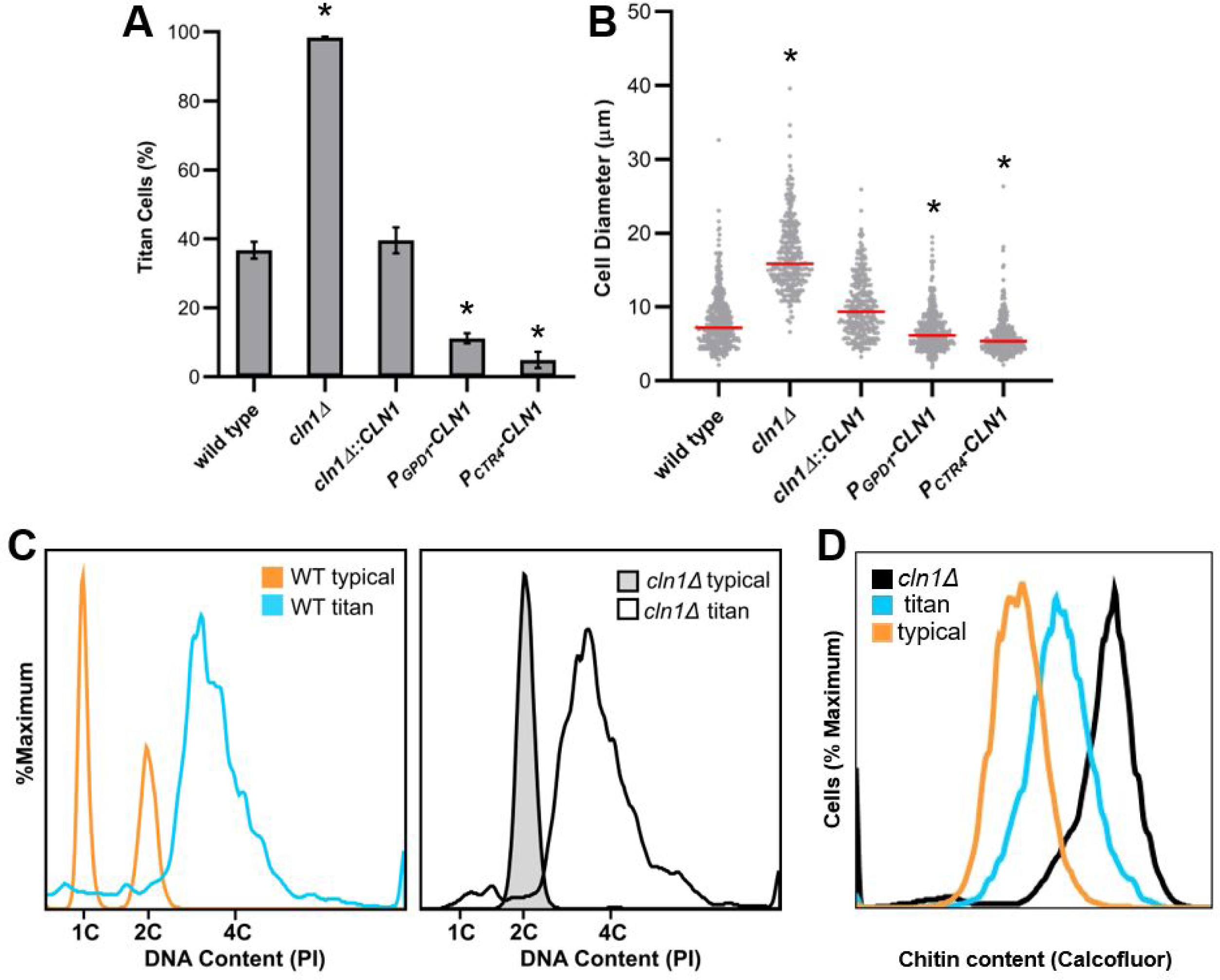
Cln1 negatively regulates *in vivo* titan cell formation. Mice were infected via inhalation with 5x10^4^ cells of the wild type strain (KN99α), the *cln1Δ* deletion, the *cln1Δ::CLN1* complement, or the over-expression strains *P_GPD1_-CLN1* or *P_CTR4_-CLN1*. Titan cell formation in the lungs was analyzed at 3 days post-infection. A) Percentage of titan cells was determined based on a cell body size threshold of 10 µm, excluding the capsule, for >300 cells per mouse. Error bars indicate SD, n ≥3 mice per strain. *p<*0.05* compared to wild type by Student’s t-test with Welch’s correction. B) Cell body size, excluding the capsule, was also plotted for >300 cells per mouse as a visual representation of the distribution of cell size in the different strains. Median cell size is indicated by the red line. *p<0.05 compared to wild type by Kruskal Wallis with Dunn post test. C) *cln1Δ* cells isolated from the lungs of mice were fixed, stained with propidium iodide, and analyzed by flow cytometry for DNA content to determine cell ploidies within the population. Wild type *in vitro* grown cells (orange), wild type titan cells (blue) and *in vitro* grown *cln1Δ* cells (gray), and a diploid strain (not shown) were used as controls. D) Wild type and *cln1Δ* cells isolated from the lungs of mice were fixed, stained with calcofluor white, and analyzed by flow cytometry to determine cell wall chitin content. The wild type population was split based on cell size into the typical (orange) and titan (blue) cell subsets for chitin analysis.

### Release from stationary phase into nutrient replete media induces morphological defects in the *cln1Δ* mutant

We showed Cln1 negatively regulates titan cell formation during infection, resulting in a dramatic increase in titan cell formation, and that the lung environment triggers an unbudded G2 arrest. Our preliminary *in vitro* studies of the *cln1Δ* deletion strain revealed only minor defects in log phase growth in nutrient replete media compared to wild type cells. Overnight cultures of the *cln1Δ* deletion strain exhibited a slight increase in overall cell size, were primarily unbudded 2C cells, and exhibited a more severe growth defect at 37°C when compared to wild type cells (**Supplementary Figures SF1 and SF3**). These results led us to hypothesize that Cln1 regulates the unbudded G2 arrest. To test this hypothesis, we took advantage of our ability to produce an unbudded 2C population through nutrient starvation *in vitro* and evaluated the capacity of *cln1Δ* cells to enter and exit stationary phase.

The ability to enter stationary phase was assessed based on cell concentration, morphology, ploidy, and viability. The *cln1Δ* deletion strain entered stationary phase at a slightly lower cell concentration than wild type, but was otherwise able to maintain an appropriate stationary population (**Figure 4A**). While the majority of the wild type cells remained unbudded throughout nutrient deprivation, buds were observed starting at day 2 in roughly 40% of the *cln1Δ* cells (**Figure 4B**). Cells in the *cln1Δ* strain had 2C ploidy under nutrient starvation at an earlier timepoint than the wild type, although it should be noted that more of the *cln1Δ* cells are in 2C in the log phase cultures at the start of the experiment (**Figure 4C**). Coincident with new bud formation on day 2, the *cln1Δ* deletion also developed a small population of 4C cells starting on day 2. Finally, no difference in viability was observed between the *cln1Δ* and wild type cells when tested using spot assays on YPD medium (**Figure 4D**). These data suggest Cln1 is not required for the unbudded G2 stationary phase arrest or viability, but may be involved in maintaining control over cell morphology and ploidy during the unbudded G2 arrest.

**Figure 4.**
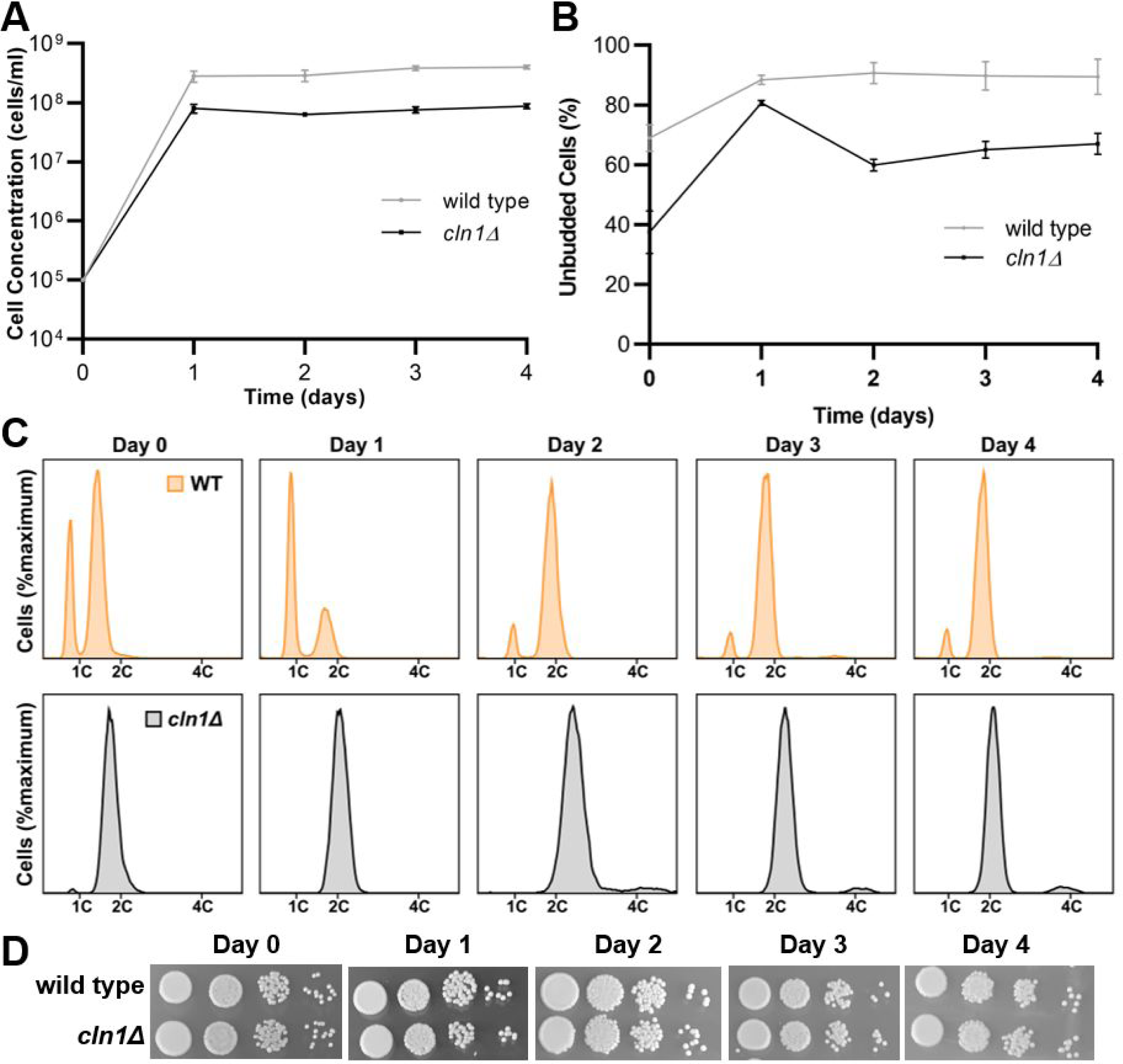
*cln1Δ* cells arrest as 2C cells and maintain viability during nutrient deprivation. Wild type and *cln1Δ* cells were grown in YPD liquid media for 4 days and assessed for entry into stationary phase at various time points based on cell concentration, morphology, viability and ploidy. Error bars represent SD from three biological replicates. A) Wild type (grey) and *cln1Δ* cells (black) displayed a plateau in cell growth after 1 day of culture. B) By 1 day, when nutrients became limited, both wild type and *cln1Δ* cells were primarily unbudded, while the *cln1Δ* cells contained more budded cells at the later time points. C) DNA content analysis showed both strains arrested as 2C cells. The *cln1Δ* cells were primarily 2C throughout the experiment, while the wild type cells arrested as 2C cells only after 2 days in YPD liquid media. D) Spot assays were performed at days 0, 1, 2, 3, and 4 of nutrient deprivation and no differences in viability were observed.

To more thoroughly assess cell morphology after stationary phase release, we released cells into liquid media and evaluated the cell morphology of each strain at various time points (**Figure 5**). After release, the *cln1Δ* cells exhibited a delay in budding and an aberrant bud morphology (**Figure 5A**). By 3 hours after release, *cln1Δ* cells had an elongated bud morphology resulting in pseudohyphae-like cells, with some daughter cells initiating another bud site without undergoing cytokinesis (**Figure 5A**). This aberrant bud morphology peaked at 5 hours after release in the *cln1Δ* mutant and was transient; by 24 hours after release, the *cln1Δ* cells consisted primarily of budded/unbudded cells, containing only a minority of cells with an aberrant morphology that was not significantly different from the wild type culture (p=0.163743) **(Figure 5B, C).** These data show Cln1 is important for regulating morphological changes associated with reentry into the cell cycle following the stationary phase unbudded G2 arrest.

**Figure 5.**
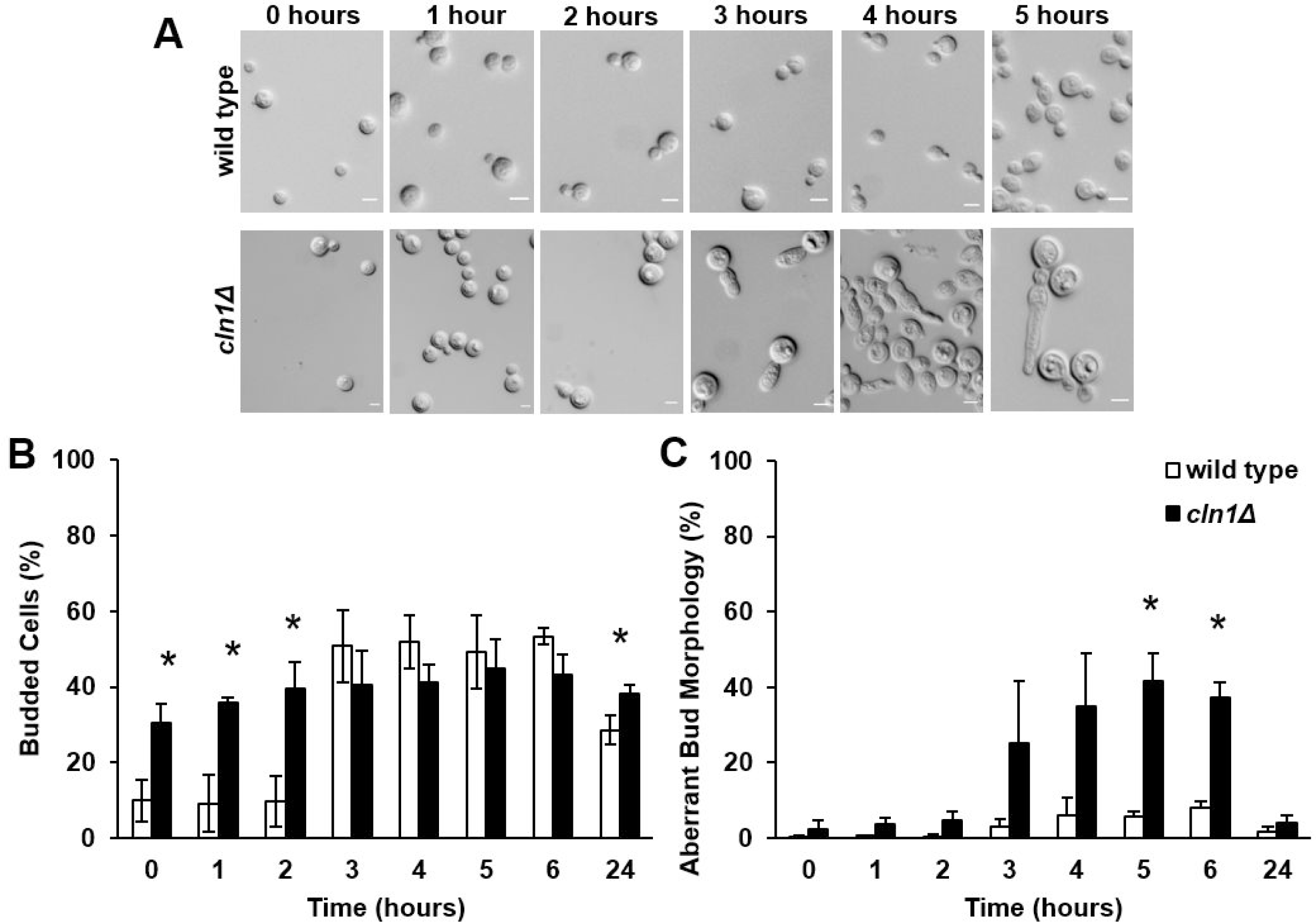
*cln1Δ* cells exhibit a transient aberrant budding morphology after release from nutrient deprivation. After 3 days incubation under nutrient starvation conditions, wild type and *cln1Δ* cells were pelleted and resuspended into fresh YPD liquid media. Error bars represent SD from three biological replicates. A) The *cln1Δ* cells exhibited an elongated bud morphological defect and delayed bud formation. Scale bar = 5 µm. B) While the wild-type cells initiated budding at 3 hours post-transfer, no increase in *cln1Δ* cell budding was observed post-transfer. C) The *cln1Δ* cells had a transient aberrant bud morphology that peaked at 5 hours post-transfer but was not observed at 24 hours. *p<0.05 by Student’s t-test with Welch’s correction.

### Cln1 binding alters Cdk1 kinase activity

We also measured the expression of *CLN1* in wild type cells after stationary phase release and the impact of Cln1 protein binding on activity of cyclin dependent kinase 1, Cdk1, the primary cyclin dependent kinase regulating cell cycle progression in *C. neoformans* (39). Expression profiling by RT-qPCR showed *CLN1* expression peaks 30 minutes after release from stationary phase (**Figure 6A**). In this analysis, the cells released from stationary phase initiated bud formation 30 minutes after release with peak budding occurring by 60 minutes after release (data not shown). To monitor interactions between Cln1 and Cdk1, we generated fusion proteins using His and Myc tags, respectively (**Supplementary Figure SF4**). Consistent with the expression data, peak interaction of Cln1-His with Cdk1-Myc occurred between 30 and 60 minutes after release from stationary phase (**Figure 6B and 6C**). This interaction increased the Cdk1 kinase activity (**Figure 6D**). Combined, these data show that peak *CLN1* expression results in Cln1 binding to Cdk1, increasing the Cdk1 kinase activity that ultimately results in initiation of bud formation.

### *C. neoformans* Cln1 contains both *S. cerevisiae* Cln and Clb cyclin motifs

*C. neoformans* Cln1 is a homolog of the *S. cerevisiae* Cdc28 cyclins (**Table 1**). In *S. cerevisiae*, there are nine Cdc28 cyclins: the G1 cyclin Cln3; two G1/S transition cyclins, Cln1 and Cln2; two S cyclins, Clb5 and Clb6; two G2/M cyclins, Clb3 and Clb4; and two M cyclins, Clb1 and Clb2. In contrast, the *Cryptococcus* species have only three Cdc28 cyclin homologs. Additionally, although both *C. neoformans* and *S. cerevisiae* divide by budding, there are key differences in their cell cycles (23, 25) (**Figures 1 and 2**). To investigate the evolutionary relationship between *C. neoformans* Cln1 and the *S. cerevisiae* Cdc28 cyclins, we performed a phylogenetic analysis and compared the conserved motifs among the Cdc28 cyclins from both *C. neoformans* and *S. cerevisiae* (**Figure 7A** **and Supplementary Figure SF5**). Interestingly, *C. neoformans* Cln1 exhibited similarity with both *S. cerevisiae* Cln and Clb cyclins and had highest motif similarity to *S. cerevisiae* Clb3 (**Figure 7A**).

**Figure 6.**
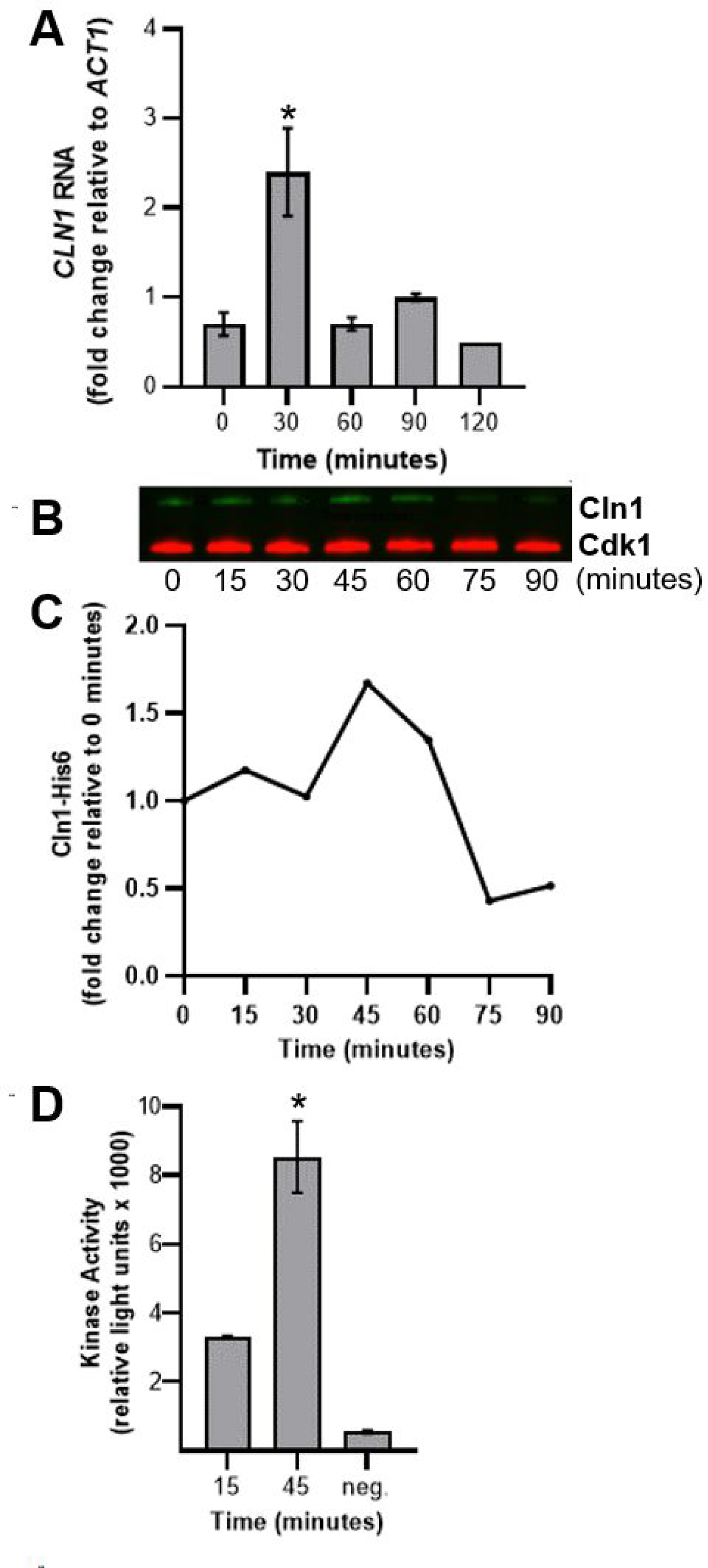
Cln1 activates Cdk1 to initiate bud formation. A) CLN1 RNA levels in wild type cells were analyzed by RT-qPCR at 30-minute intervals after release from stationary phase into nutrient replete media. Expression levels were normalized to the ACT1 housekeeping gene to determine changes in gene expression. Error bars represent SD of the Ct values of 3 technical replicates. Data are representative of 3 biological replicates. *p<0.05 compared to t=0 by Student’s t-test with Welch’s correction. B-C) Cells were harvested at 15-minute intervals and the amount of Cln1-His6 co-precipitated with Cdk1-Myc was determined at each time point by western blot (B) and then quantified based on the 0 minute time point (C). D) Cdk1 kinase activity was determined at 15 and 45 minutes. The negative control consisted of the 1x kinase buffer only. Error bars represent SD of 3 technical replicates. *p<0.05 compared to t=15 by Student’s t-test.

**Figure 7.**
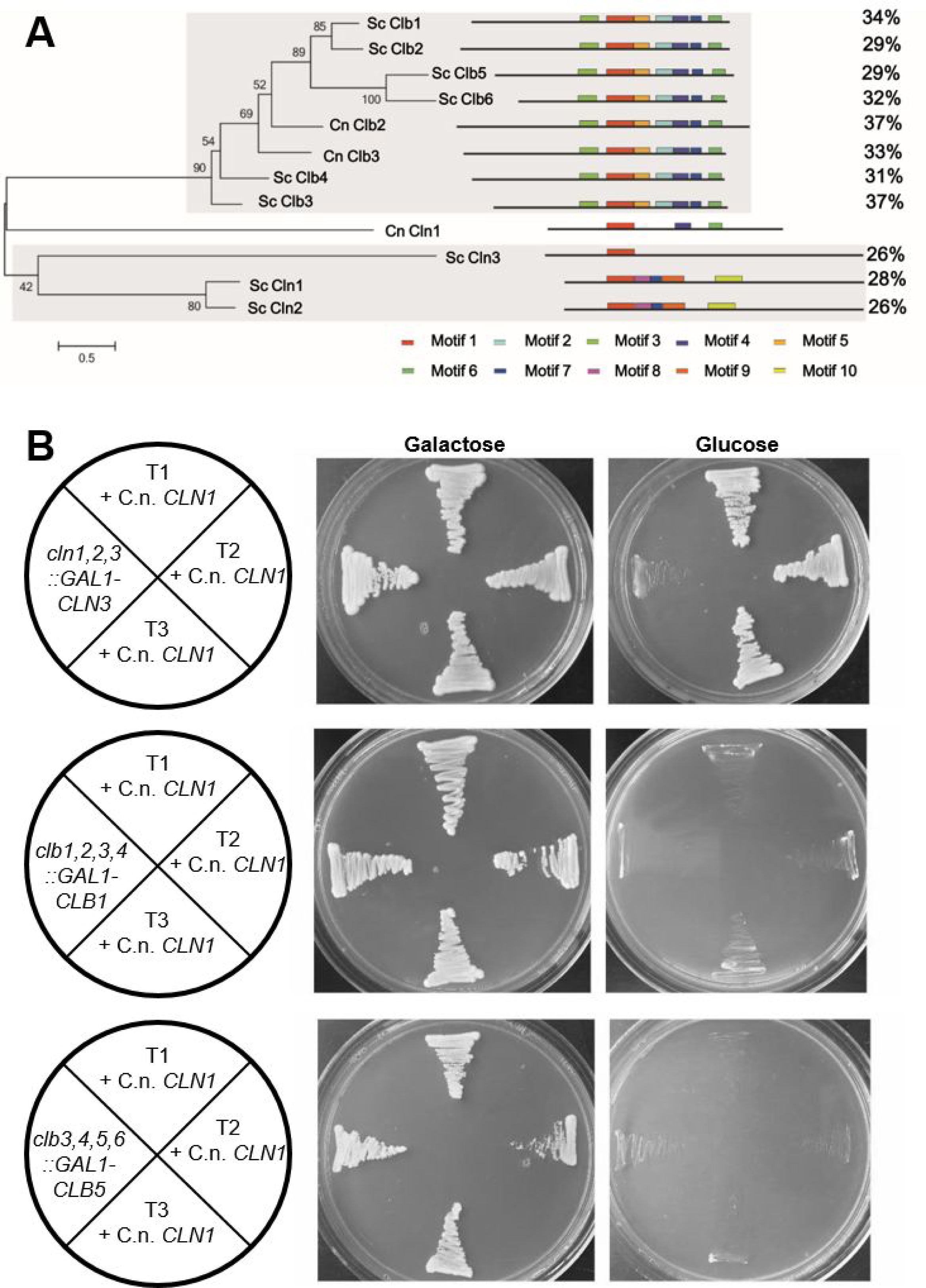
Motif analysis of *C. neoformans* Cln1 shows it contains both *S. cerevisiae* Cln and Clb motifs but only complements Cln cyclins in *S. cerevisiae*. A) Phylogenetic analysis based on protein motifs of the Cdc28 family of cyclins from *C. neoformans* (Cln1, Clb2 and Clb3) and the *S. cerevisiae* (Cln1, Cln2, Cln3, Clb1, Clb2, Clb3, Clb4, Clb5, and Clb6) showed *C. neoformans* Cln1 had highest homology to the *S. cerevisiae* Clb proteins. B) *C. neoformans CLN1* expressed in *S. cerevisiae* was able to rescue growth of a *cln1,2,3* triple but not *clb1,2,3,4* or *clb3,4,5,6* quadruple mutants. *C. neoformans CLN1* was expressed with a constitutive *S. cerevisiae* promoter and all *cln1,2,3::GAL-CLN3* + *C. neoformans CLN1* transformants (T1, T2, or T3 + C.n. *CLN1*) produced growth on the non-permissive glucose medium whereas none of the *clb1,2,3,4::GAL1-CLB1* + *C. neoformans CLN1* or *clb3,4,5,6::GAL-CLB5* + *C. neoformans CLN1* transformants had growth on the non-permissive glucose medium.

The *C. deneoformans CLN1* gene was previously shown to complement the *S. cerevisiae* Cln cyclins (40), but was not tested for Clb cyclin complementation. In *C. neoformans*, peak *CLN1* expression occurs after release from stationary phase at the G2 to M transition (**Supplementary Figure SF1**). In addition, the elongated bud morphology observed in the *cln1Δ* cells is similar to the elongated bud phenotype observed in *S. cerevisiae* cells deficient in the mitotic cyclin Clb2 that regulates the G2 to M cell cycle transition (41, 42). These data led us to hypothesize that the Cln1 cyclin in *C. neoformans* may act at the G2 to M transition in the unbudded G2 arrest stress cell cycle, and given the motif similarity may be able to also complement the *S. cerevisiae* Clb cyclins.

To test this hypothesis, we analyzed whether *C. neoformans* Cln1 can functionally complement the *S. cerevisiae* Cln and Clb cyclins. The *S. cerevisiae cln1Δ cln2Δ cln3Δ* triple mutant when complemented with *CLN3* fused to a galactose inducible promoter is deficient for Cln cyclins in the presence of glucose, resulting in no growth. In contrast, the triple mutant strain grows on galactose where *GAL-CLN3* is expressed. Similar to *C. deneoformans* (40), addition of the *C. neoformans CLN1* gene coding region under control of the *S. cerevisiae GPD1* constitutive promoter restored growth of the *S. cerevisiae cln1Δ cln2Δ cln3Δ GAL-CLN3* cells on glucose, showing that *C. neoformans* Cln1 can functionally complement the *S. cerevisiae* Cln cyclins (**Figure 7B**). When we performed the same complementation experiment with the Clb deficient strains *clb1Δ clb2Δ clb3Δ clb4Δ GAL-CLB1* or *clb3Δ clb4Δ clb5Δ clb6Δ GAL-CLB5*, the *C. neoformans* Cln1 was unable to functionally complement the *S. cerevisiae* Clb cyclins (**Figure 7B**). Taken together, these comparative studies between the *S. cerevisiae* cyclins and *C. neoformans* Cln1 showed that Cln1 has homology to both the Cln and Clb cyclins in *S. cerevisiae* but only functionally complements the *S. cerevisiae* Cln cyclins.

### Low Cln1 expression after unbudded G2 arrest leads to polyploid titan cell formation

The functional complementation experiments in *S. cerevisiae* only identified a possible role for *C. neoformans* Cln1 in regulating early phases of the cell cycle that are controlled by Cln cyclins in *S. cerevisiae*. Yet our observations that *C. neoformans* Cln1 expression peaked upon initiation of budding after release from stationary phase growth arrest, the elongated bud morphology observed in the *cln1Δ* deletion upon release from stationary phase *in vitro*, the homology to the *S. cerevisiae* Clb cyclins, and activation of the *C. neoformans* cyclin-dependent kinase Cdk1 by Cln1 at bud formation all indicated a role for *C. neoformans* Cln1 in later phases of the cell cycle. Given the further observations that *C. neoformans* has a stress-induced unbudded G2 arrest, we hypothesized that Cln1 is critical for maintaining the balance between cell growth and DNA replication during this unbudded G2 arrest.

We first explored the association between *CLN1* expression levels in titan and typical sized cells *in vivo*, testing the hypothesis that low Cln1 levels would be unable to maintain control of the unbudded G2 arrest and result in titan cell formation. RT-qPCR analysis of *CLN1* RNA isolated from typical and titan cells and normalized to cell number showed lower *CLN1* expression in titan cells compared to typical sized cells *in vivo* (**Figure 8A**, p = 0.0004). While these results provide indirect support for our hypothesis that low levels of Cln1 in a cell would be unable to maintain the unbudded G2 arrest and lead to titan cell formation, we were unable to adequately manipulate the *in vivo* system.

**Figure 8.**
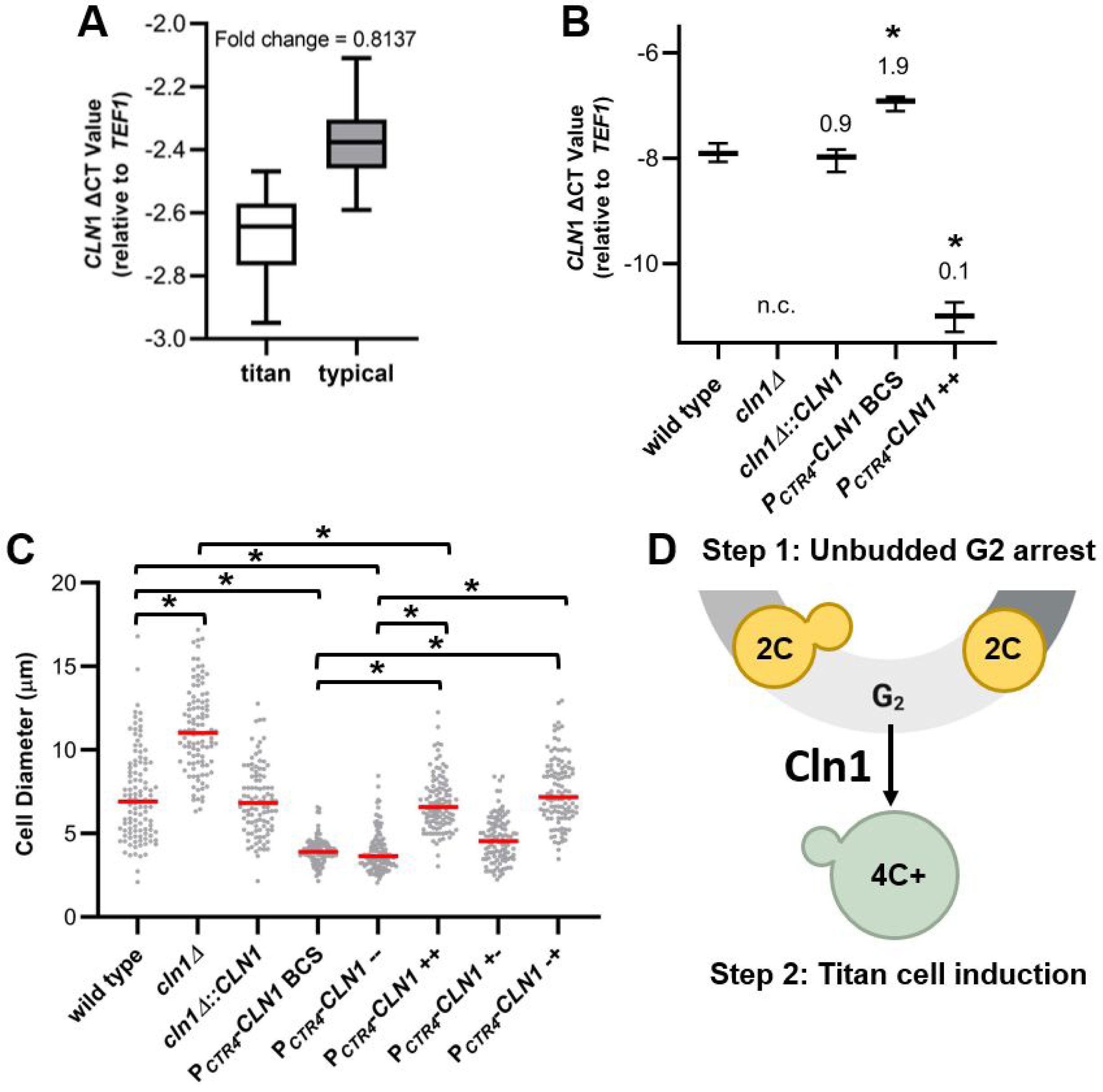
Low *CLN1* expression after unbudded G2 arrest induces titan cell formation. A) RT-qPCR showed higher *CLN1* levels in typical size cells compared to titan cells isolated from the lungs of mice. Expression levels were normalized to both cell number and the TEF1 housekeeping gene to determine change in cycle threshold (ΔCT). Fold decrease in gene expression in titan compared to typical cells was calculated using 2-ΔΔCt. Data presented are the ΔCT values from three biological replicates. p = 0.0004 by Student’s t-test. B) RT-qPCR analysis of CLN1 expression *in vitro* in the wild type, *cln1Δ*, *cln1Δ::CLN1*, and when the copper repressible promoter strain was grown with 400 µM of the copper chelator BCS (*P_CTR4_-CLN1* BCS) or 25 µM copper (P_CTR4_-CLN1++). No expression change was detected in the *cln1Δ* strain (n.c). Fold change compared to wild type is indicated above each bar. Data presented are the ΔCT values from three technical replicates. *p ≤ 0.0015 compared to wild type by one-way ANOVA with a Dunnett correction for multiple comparisons. C) *In vitro* titan cell formation with wild type, *cln1Δ*, *cln1Δ::CLN1*, and *P_CTR4_-CLN1* showed *CLN1* expression only affects titan cell formation after unbudded 2C cells are already formed. *In vitro* titan cell formation was induced using an initial stationary phase culture to induce unbudded G2 arrest followed by a second incubation under hypoxic conditions to induce titan cell formation (Hommel et al., 2018). CLN1 expression was manipulated using the P_CTR4_-CLN1 strain and copper addition during the initial stationary culture (+-), second incubation (-+), both (++), or neither (--). The cell diameter of at least 100 cells was measured at the end of the protocol to determine *in vitro* titan cell formation. Median cell size is indicated by the red line. *p < 0.0001 by Kruskal-Wallis test with Dunn’s post test correction. D) Our results show Cln1 is necessary in the “stress” cell cycle after the unbudded 2C cells are already formed. Loss or reduction of *CLN1* expression in cells after unbudded G2 arrest results in polyploid titan cells (green cells) only with concomitant environmental stimuli.

Methods to generate titan-like cells *in vitro* were recently identified (43–45), allowing us to manipulate *CLN1* expression (**Figure 8B**) and monitor the effect on titan cell formation. Furthermore, the *in vitro* titan cell assays require a two-step process to induce titan cell formation. The first step induces the unbudded G2 arrest and the second step induces titan cell formation using environmental cues. By manipulating *CLN1* expression during these two steps, we can pinpoint whether Cln1 is critical to maintain the unbudded G2 arrest, to control titan cell formation, or is acting at both steps in the process of titan cell formation.

As previously shown for *in vivo* titan cell formation, the *cln1Δ* deletion strain had increased *in vitro* titan cell formation compared to the wild type strain (p<0.0001) and no *CLN1* gene expression (**Figure 8B and 8C**). The *P_CTR4_-CLN1* strain had decreased titan cell formation compared to the wild type strain in the *in vitro* assay, both with and without depletion of copper in the media using BCS (p<*0.0001*), that resulted in an almost a 2-fold increase in *CLN1* expression compared to wild type (**Figure 8B and 8C**). When copper was present in the media (*P_CTR4_-CLN1* ++), the *CLN1* expression in the *P_CTR4_-CLN1* strain decreased significantly (fold change in expression = 0.1) and resulted in similar titan cell formation as the wild type strain (p>0.9999) but lower than the *cln1Δ* deletion strain (p<0.0001) (**Figure 8B and 8C**). The observation that the *P_CTR4_-CLN1++* strain in the presence of copper does not exhibit the same increase in titan cell formation as the *cln1Δ* deletion strain has two possible explanations. The most likely explanation is that the copper repressible promoter is “leaky” and there is still a low level of *CLN1* expression. Consistent with this explanation, the P_CTR4_-CLN1 strain in the presence of copper does not display complete abolishment of *CLN1* expression (**Figure 8B**). A second explanation is that some of the signals leading to titan cell formation are present in the *CLN1* promoter, and replacement of the *CLN1* promoter with the *CTR4* promoter produced a strain unable to overproduce titan cells when *CLN1* is overexpressed.

The conditions we used for *in vitro* titan cell formation required an initial first step that induced the unbudded G2 arrest using nutrient starvation prior to shifting the cells to titan inducing conditions in the second step. Thus, we dissected the role of Cln1 in titan cell formation using the *P_CTR4_-CLN1* strain both in the initial incubation that results in the G2 arrest and the subsequent incubation that produces the titan cells (**Figure 8C**). When we repressed *CLN1* only in the initial incubation (*P_CTR4_-CLN1* -+) to test if Cln1 was required during the G2 arrest, we observed no differences in titan cell formation compared to the *P_CTR4_-CLN1 BSC and P_CTR4_-CLN1--* controls (p>0.9999). In contrast, when *CLN1* was repressed in the second incubation *(P_CTR4_-CLN1+-)*, titan cell formation increased compared to the *P_CTR4_-CLN1 BSC and P_CTR4_-CLN1--* controls (p<0.0001). These results show Cln1 activity is only necessary after the unbudded G2 arrest. Under appropriate environmental conditions, insufficient *CLN1* expression results in polyploid titan cell formation following the unbudded G2 arrest (**Figure 8D**).

## Discussion

Although titan cells play a role in virulence in the pathogenic yeast *C. neoformans*, the mechanisms underlying titan cell formation are still largely unknown. Previous studies exploring the host signals and signal transduction pathways stimulating titan cell formation showed their production is tightly regulated and has similarities to morphological switches in other fungal species that are cell cycle regulated (43, 46–50). We characterized the cell cycle of *C. neoformans* during *in vivo* and *in vitro* stress to better understand potential mechanisms leading to titan cell formation and linked these studies to identification of the putative cyclins and CDKs to investigate their roles in titan cell formation. Our results show that titan cell formation is cell cycle regulated and the cyclin Cln1 negatively regulates titan cell formation. Additionally, we show that *C. neoformans* produces unbudded G2 cells in response to *in vivo* and *in vitro* stresses and that Cln1 is critical for regulation of morphogenesis after this G2 arrest, leading us to postulate that the unbudded 2C cell is a precursor to titan cells (**Figure 8D**).

Previous studies identified Cln1 in *Cryptococcus*, but the link to titan cell formation, and its exact role in the cell cycle remained unclear (40, 51, 52). To better understand these mechanisms and the role of Cln1 in the unbudded G2 arrest, we assessed the ability of *C. neoformans* Cln1 to functionally complement Cln and Clb cyclins in *S. cerevisiae*. Cln1 in the closely related species *C. deneoformans* had already been shown to complement the *S. cerevisiae* Cln cyclins (40), so we were unsurprised that the *C. neoformans* Cln1 was also able to complement the Cln cyclins. Additionally, we showed that *C. neoformans* Cln1 does not functionally complement the *S. cerevisiae* Clb cyclins. This result may seem contradictory to our data showing that *C. neoformans* Cln1 is involved in cell cycle regulation after the unbudded G2 arrest. However, there are a number of important differences between the *S. cerevisiae* and *C. neoformans* cell cycles that may shed light on the unique functionality of Cln1 in *C. neoformans*.

First, in *S. cerevisiae*, the Cln cyclins are associated with the initiation of START, a cell cycle commitment point that occurs at the G1-S transition. The presence of START in *S. cerevisiae* ensures budding and DNA replication are tightly coordinated (53). In contrast, our cell sorting studies in *C. neoformans* show that DNA replication without bud formation readily occurs both *in vivo* and *in vitro*, and are consistent with previous microscopy studies in both *C. neoformans* and *C. deneoformans* (18, 26–29, 54). These data suggest that START does not exist in *Cryptococcus.* This lack of START likely allows Cln1 to be multifunctional across the *C. neoformans* cell cycle.

An alternate possibility is that *C. neoformans* Cln1 has novel functions in the later part of the cell cycle due to its homology to the Clb cyclins. The elongated bud morphology observed in the *cln1Δ* cells is similar to the elongated bud phenotype observed in *S. cerevisiae* cells deficient in the mitotic cyclin Clb2 that regulates the G2 to M cell cycle transition and inhibits polarized bud growth to initiate the apical to isotropic switch that allows the cell to maintain a normal bud morphology. Clb2-deficient cells in *S. cerevisiae* exhibit a G2 mitotic delay, characterized by elongated hyperpolarized buds (41, 55, 56). It should be noted, however, that the elongated bud phenotype is only transiently observed in the *C. neoformans cln1Δ* mutant *in vitro* upon release from stationary phase; and a similar elongated bud phenotype is also seen in *S. cerevisiae* when *CLN1* is expressed during G2 (41).

Finally, the *C. neoformans* cell cycle shares more similarities to that of mammals and other multicellular eukaryotes than the other model yeasts, such as *S. cerevisiae* and *S. pombe*. For example, *C. neoformans* undergoes a stepwise kinetochore assembly (25) and contains several microtubule organizing centers (MTOCs) (57). In addition, while *S. cerevisiae* and *S. pombe* have a closed mitosis (58–60), *C. neoformans* has a semi-open mitosis (25) that is more similar to the open mitosis observed in mammals. In addition, many multicellular eukaryotes, including humans, also have cell cycles that do not rely heavily on START; this cell cycle commitment checkpoint is often referred to as the “restriction point” in higher eukaryotes. Specifically, cancer cells are well known for their ability to bypass the restriction point, often resulting in polyploidy (61).

Our studies show that *CLN1* expression increases upon release from stationary phase and is also lower in titan cells compared to typical sized cells *in vivo*. During log phase growth *C. neoformans* utilizes a cell cycle characterized by synchronous DNA replication and bud formation that is also observed in *S. cerevisiae* (25). During stationary phase and *in vivo* growth, we observed a pronounced G2 arrest. This unbudded G2 arrest was observed previously in various C. *deneoformans* and *C. neoformans* strains in response to other *in vitro* stresses in addition to nutrient deprivation (26, 27), including high temperature (29) and hypoxia (28), and thus we refer to this alternative cell cycle as the *Cryptococcus* “stress cell cycle”. Our *in vitro* stationary phase and titan cell formation experiments with the *P_CTR4_-CLN1* strain are important for several reasons. First, the stationary phase studies show Cln1 has minimal effect on entry into the G2 arrest and are corroborated by the titan cell formation experiments. Second, because *in vitro* titan cell formation utilizes a two-step process we can pinpoint where Cln1 acts in the cell cycle to regulate titan cell formation. Our studies revealed that Cln1 functions after the G2 arrest – at the critical decision point when the cell either remains arrested, undergoes DNA re-replication to form a titan cell, or re-enters the cell cycle to produce a daughter cell.

In addition, the *in vitro* titan cell formation experiments highlight the fact that while titan cell formation is affected by low Cln1 levels in the cell, low Cln1 is not the only mechanism underlying titan cell formation. Under conditions that do not promote titan cell production, such as *in vitro* log and stationary phase growth, the *cln1Δ* mutant does not produce titan cells. These data show *C. neoformans* needs additional signals to produce titan cells and is consistent with previous studies showing that multiple signal transduction pathways affect titan cell morphogenesis and that production of titan cells *in vitro* requires complex environmental conditions that involve multiple forms of induction (37). There are a variety of signals that affect titan cell formation, but our data highlight the need for the unbudded 2C cell as one of the earliest of these required signals.

The mechanism by which low Cln1 levels in the cell lead to titan cell formation still remains unclear. We postulate that low Cln1 levels in the cell result in DNA replication in the absence of cell division, or DNA re-replication, under environmental conditions that stimulate isotropic growth of the cell, ultimately producing the titan cell phenotype (**Figure 8D**). This DNA re-replication may result from premature mitotic exit, a process referred to as mitotic slippage. Mitotic slippage is known to form polyploid cancer cells (62, 63). Alternatively, Cln1 deficient cells may not arrest in G2 but instead continue to undergo mitosis – resulting in nuclear division in the absence of cell division, possibly leading to polyploidy through mitotic collapse such as that previously described in *C. albicans* (64). In *C. albicans* mitotic collapse is associated with the production of a trimera cell morphology. In our studies we did not observe trimeras or evidence of mitotic spindle collapse in our time lapse microscopy (Altamirano, Fu, Nielsen, unpublished data). However, in *C. neoformans* lack of START and the unbudded G2 arrest may make the processes of mitotic slippage and mitotic collapse morphologically indistinguishable.

Cell cycle arrest is an important stress response mechanism in many organisms (65, 66). Both our *in vivo* studies and the previous *in vitro* studies with nutrient, hypoxia, and temperature stress identified the production of 2C unbudded cells in response to various stresses (26–29, 54). Why might 2C arrest be beneficial under stressful conditions? Cell cycle arrest could promote changes in cell morphology or development that allows for increased adaptation to external stress signals. For example, Fu et al. previously described the ability of *C. neoformans* to undergo hyphal formation and monokaryotic fruiting after an unbudded 2C arrest. It is possible that *C. neoformans* incorporated the cell cycle arrest into a global stress response that ultimately evolved into generation of the titan cell phenotype. Not unsurprisingly, the titan cell phenotype includes traditional aspects of cell cycle regulation such as ploidy alterations, including ploidy increases during titan cell formation and ploidy reductions during titan cell division, and morphological alterations such as isotropic growth. However, the titan cell phenotype also includes changes linked to defense strategies such as alterations in cell wall, melanin, and capsule structure (19, 22). Cln1 was previously linked to these cell surface phenotypes (51, 52), with the association likely related to the role of Cln1 in the stress cell cycle and its role in cell defense. In support of this conclusion, Garcia-Rodas (51) used the *Galleria* model to bypass the high temperature growth defect of the *cln1Δ* mutant and showed that the mutant had lower survival inside *Galleria* hemocytes at 30°C, highlighting a critical role for the stress cell cycle and cellular defenses *in vivo*.

Finally, our studies show the unbudded G2 arrest and stress cell cycle that gives rise to the polyploid titan cells and subsequent aneuploid daughter cells with novel traits is a pre-programmed response to stress in *C. neoformans* (20). Given the similarities between the *C. neoformans* cell cycle and those of multicellular eukaryotes, titan cell formation and aneuploid daughter cell production may be an excellent system to understand the fundamental cellular processes that underlie eukaryotic polyploid cells. The ability to study the cause and consequence of ploidy increases and reductions in a natural, non-engineered, single celled organism as a model for humans or higher eukaryotes promises immense future research potential.

## Materials and Methods

### Ethics statement

All mice were handled in strict accordance with good animal practice, as defined by the relevant national and/or local animal welfare bodies. All animal work was approved by the University of Minnesota Institutional Animal Care and Use Committee (IACUC) under protocol no. 1308A30852.

### Strains and culture conditions

The *C. neoformans* and *S. cerevisiae* strains used in this study are listed in Supplementary Table ST2. KN99α was used as the wild type strain unless otherwise indicated. Strains were stored in 30% glycerol at -80°C. *C. neoformans* strains were grown at 30°C in yeast extract-peptone-dextrose (YPD) agar or liquid medium (BD Biosciences, Sparks, MD). *S. cerevisiae* strains were grown at 30°C in yeast extract-peptone-galactose (YPG) agar or liquid medium. Drop-out mix synthetic minus uracil or adenine (US Biological, Salem, MA) with galactose agar was used for *S. cerevisiae* transformation. YPD agar containing 200 µg/ml nourseothricin (Sigma Aldrich, St. Louis, MO), 100 µg/ml G-418 (AG Scientific, San Diego, CA) and/or 300 µg/ml hygromycin B (EMD Millipore, Billerica, MA) was used for *C. neoformans* transformation. Mating was performed on V8 juice agar (67).

### Identification of putative cyclins and cyclin-dependent kinases in *C. neoformans*

To identify the putative cyclins in *C. neoformans*, the protein sequences of *S. cerevisiae* cyclins (Cln1, Cln2, Cln3, Clb1, Clb2, Clb3, Clb4, Clb5, Clb6, Pcl1, Pcl2, Pcl5, Pcl6, Pcl7, Pcl8, Pcl9, Pcl10, Pho80, Clg1, Ccl1, and Ssn8) and *S. cerevisiae* CDKs (Cdc28, Pho85, Kin28, Ssn3, and Ctk1) were subjected to Blastp searches against *C. neoformans* var. *grubii* H99 database from the Broad Institute with E-value cutoff set as

1. A second round of Blastp searches against the *C. neoformans* var. *grubii* H99 database was carried out using the protein sequences of the putative *C. neoformans* cyclins and CDKs with E-value cutoff set as 1.

### Strain construction

Primers used in this study are listed in Supplementary Table ST3. The deletion strains were generated by gene disruption as previously described (68). Briefly, the coding region of the putative cyclin or CDK gene was replaced by the nourseothricin N-acetyl transferase (*NAT)* drug resistance cassette. An overlap PCR product was created with the 5’ flanking region (∼1 kb), *NAT* resistance cassette, and 3’ flanking region (∼1 kb) (68). The PCR product was introduced into the KN99α strain by biolistic transformation (69). The resulting deletion mutants generated via homologous recombination were confirmed by PCR, sequencing of the PCR product, and Southern blot. For complementation of the *cln1Δ* deletion strain, an overlap PCR product was generated that contained the KN99α gene promoter, open reading frame, and terminator, a neomycin (*NEO*) drug resistance cassette (70) and the 3’ flanking region (∼1 kb). The PCR product was introduced into the mutant by biolistic transformation. The resulting complement strain generated via homologous recombination was confirmed by PCR, RT-qPCR, and phenotype analysis.

For generation of the overexpression strains, expression of *CLN1* was driven under the constitutively expressed promoter *GPD1* (33) or copper repressible promoter *CTR4* (*34*). An overlap PCR product was generated with 5’ flanking regions (∼1 kb), *NAT* resistance cassette, *GPD1* promoter (0.9 kb) or *CTR4* promoter (2 kb), and 1 kb of the coding region of the KN99α *CLN1* gene (68). The PCR product was introduced into the KN99α strain by biolistic transformation (69). The resulting overexpression strain generated via homologous recombination was confirmed by PCR and RT-qPCR under appropriate expression conditions.

The strain expressing fluorescently labelled *TUB1-GFP* under the endogenous *TUB1* promoter was constructed by first inserting *GFP-TUB1-NEO* with the native TUB1 promoter at the TUB1 locus into the *C. neoformans* wild type strain, KN99α. To tag Tub1 with GFP at the N-terminus, the *TUB1* promoter region and *TUB1* gene were first amplified by PCR from *C. neoformans* KN99α genomic DNA, and the GFP gene was amplified from pAcGFP1 (Clontech, Mountain View, CA). The three PCR fragments were fused together using overlap PCR (68). The overlap PCR product was cloned into pSC-B ampkan (Agilent Technologies, La Jolla, CA) to generate pScGFPTUB1 using a Stratagene blunt PCR cloning kit (Agilent Technologies). The neomycin (*NEO*) drug resistance marker was amplified from pJAF12 and cloned into pSC-B ampkan to make pScNEO. Using the restriction enzyme, SpeI, the *NEO* resistance gene was removed from pScNEO and subcloned into pScGFPTUB1 in the SpeI site to create the plasmid pScGFPTUB1-NEO. This plasmid was linearized using the restriction enzyme, StuI, and biolistically transformed into KN99α (Toffaletti et al., 1993). The transformants were selected on YPD agar supplemented with 100 µg/ml neomycin. Positive transformants were screened by PCR and fluorescent microscopy.

The *GFP-TUB1-NEO* insertion resulted in two copies of *TUB1* in the KN99α genome. To delete one copy of *TUB1*, disruption cassettes with nourseothricin (*NAT*) drug resistance markers flanked by ∼1 kb upstream and downstream regions of the *TUB1* gene were constructed. The *TUB1* upstream and downstream regions were amplified from the *GFP-TUB1* strain generated above, and the *NAT* drug resistance gene was amplified from plasmid pNATSTM125. Three PCR fragments were then combined using overlap PCR. The overlap PCR product was transformed into the *GFP-TUB1* strain using biolistic transformation. Transformants were selected on YPD agar supplemented with 200 µg/ml nourseothricin. Positive transformants were screened by PCR, and qRT-PCR was performed to confirm that one copy of TUB1 was deleted by comparing the *TUB1* transcript level with the KN99α parental strain (1 copy) and the *GFP-TUB1* strain prior to transformation (2 copies).

Two strains SL306 (KN99α with *NOP1-mCherry::NEO*) and SL321 (KN99a with *GFP-NOP1::NAT*) were used to generate a KN99a strain containing *NOP1-mCherry::NEO* strain by mating (71). Mating was performed on V8 juice agar (pH 7.0) at room temperature in the dark. After two weeks, individual basidiospores were micromanipulated onto YPD agar. Strains containing *NOP1-mCherry::NEO* were identified using fluorescence microscopy. A mating type a version of the *Nop1-mCherry::NEO* strain was identified by co-culturing with KN99a and KN99α tester strains, as described above for mating.

The *NOP1-mCherry::NEO GFP-TUB1::NAT* strain was generated by mating the KN99a *NOP1-mCherry::NEO* with the KN99α *GFP-TUB1::NAT* strain. Single spores were isolated by microdissection, cultured on YPD agar containing both NAT and NEO, and the mating type of the resulting progeny were determined as described above. Progeny strains were analyzed by fluorescence microscopy to identify strains with both GFP-labeled microtubules and mCherry-labeled nucleolus.

To generate the *CLN1-His*_6_ strain, an overlap PCR product was created with the *CLN1* ORF with a His tag at the 3’ end (∼1 kb), *GPD1* terminator (∼0.3 kb), *NEO* resistance cassette, and 3’ flanking regions (∼1 kb). The PCR product was introduced into the KN99α strain by biolistic transformation. To generate the *CDK1-Myc* strain, an overlap PCR product was created with the *CDK1* ORF with the Myc tag at the 3’ end (∼1 kb), *GPD1* terminator (∼0.3 kb), *NAT* resistance cassette, and 3’ flanking regions (∼1 kb). The PCR product was introduced into the KN99α strain by biolistic transformation. The resulting deletion mutants generated via homologous recombination were confirmed by PCR and sequencing of the PCR product. To generate the *CLN1-His+ CDK1-Myc* strain, a second round of biolistic transformation was carried out by introducing the overlap PCR product of the *CLN1-His*_6_ into the *CDK1-Myc* strain.

### Confirmation of the essential cyclins in *C. neoformans*

The diploid strain KN99α/KN99a was created as described previously (72) using KN99α *NOP1-mCherry::NEO* and KN99a *14-3-3-GFP::HYG* as the progenitor strains. To generate the KN99a *14-3-3-GFP::HYG* strain, an overlap PCR product was created with the *14-3-3* ORF (∼1 kb), GFP, *GPD1* terminator (∼0.3 kb), *NEO* resistance cassette, and 3’ flanking regions (∼1 kb). The PCR product was introduced into the KN99a strain by biolistic transformation. The resulting strain generated via homologous recombination was confirmed by PCR and sequencing of the PCR product. A co-culture of KN99α *NOP1-mCherry::NEO* and KN99a *14-3-3-GFP::HYG* cells were incubated on V8 juice agar overnight at 25°C in the dark. The co-culture was transferred to YPD agar containing *NEO* and HYG to select the diploid strain KN99α *NOP1-mCherry::NEO*/KN99a *14-3-3-GFP::HYG* at 37 °C. The overlap PCR product for the putative essential gene containing the 5’ flanking regions (∼1 kb), *NAT* resistance cassette, and 3’ flanking regions (∼1 kb) was then introduced into the diploid strain by biolistic transformation as described above. The resulting transformants were selected on YPD agar containing NAT, NEO, and HYG at 37°C. The diploid deletion strains were screened by PCR to confirm that only one copy of the candidate gene was disrupted by *NAT*. The occurrence of double deletion diploid strains was used as evidence that the gene was not essential. Single deletion diploid strains were then sporulated on V8 juice agar at 25°C in the dark. One hundred basidiospores were microdissected onto YPD agar and the resulting colonies were screened for *NAT* resistance. Failure to obtain *NAT* resistant colonies was considered indicative that spores containing the putative essential gene deletion construct were inviable, and these genes are essential in haploid *C. neoformans*.

### Titan cell formation *in vivo*

Cells were grown overnight in YPD liquid medium at 30°C, then washed three times with PBS and resuspended in phosphate buffered saline (PBS) at a concentration of 1×10^6^ cells/ml based on hemocytometer count. Groups of 6- to 8-week-old female A/J mice (Jackson Labs, Bar Harbor, ME) were anesthetized by intraperitoneal pentobarbital injection. Four mice per treatment were infected intranasally with 5×10^4^ or 5×10^6^ cells in 50 µl PBS. Infected mice were sacrificed at 3 or 14 days post-infection by CO_2_ inhalation. Lungs were removed and then homogenized in 20 ml of Hanks’ balanced salt solution (HBSS) with 0.8 mg/ml collagenase type I (Life Technology, Grand Island, NY). The cell suspension was incubated for 1 hour at 37°C with continuous shaking then washed three times with 0.05% SDS to lyse the mammalian cells. The resulting *C. neoformans* cells were fixed in 3.7% formaldehyde, pelleted, and resuspended in PBS. At least 100 *C. neoformans* cells per mouse were analyzed for cell body size by microscopy. Cells were classified as typical sized cells (<10 μm in cell body diameter excluding the capsule) or titan cells (>10 μm in cell body diameter excluding the capsule).

### Cell synchronization

Cells were grown in 50 ml YPD liquid medium in a 250 ml flask at 30°C, 250 rpm, for 1.5 days to reach stationary phase. The resulting cells were diluted 1:70 with fresh YPD, again as a 50 ml YPD culture in 250 ml flask, at 30°C, 250 rpm, and samples were analyzed at 0 min, 15 min, 30 min, 45 min, 60 min, 75 min, 90 min, 105 min and 120 min for bud size analysis, flow cytometry and RT-qPCR.

### Growth curve

Cells were grown overnight in YPD liquid medium at 30°C, diluted into fresh YPD at a concentration of 5x10^4^ cells/ml, 200 μl pipetted in replicates of 3 into a microplate (tissue culture test plate 96F, TPP, Switzerland), and growth analyzed using a Synergy™ H1 Hybrid Multi-Mode Microplate Reader (BioTek Instruments, Winooski, VT). Measurements were made every 5 minutes for 75 minutes with continuous orbital shaking at 30°C and 37°C. Shaking speed was set to slow and frequency set to 282 (3 mm amplitude). Cell growth was assessed by 600 nm absorbance light scatter measurements. Data was collected using Gen5™ Data Analysis Software (BioTek Instruments, Winooski, VT).

### Flow cytometry

Propidium iodide staining was used to assess the ploidy of cells. Briefly, cells were washed and suspended in 100 µl of sterile water. Cells were then fixed with 70% ethanol, incubated at 24°C for 1 hour, and incubated at 4°C overnight. Cells were then washed and resuspended in 100 µl of RNAse A buffer (0.2M Tris pH 7.5, 20 mM EDTA) and incubated with 1 µl of RNAse A (10 mg/ml) at 37°C for 4 hours. After RNAse A treatment, cells were washed and resuspended in 900 µl of PBS. 100 µl of propidium iodide (PI) (.05 mg/ml) was added to 900 µl of cells and incubated in the dark for 30 minutes. Immediately prior to analysis, cells were sonicated at 20% amplitude for 5 seconds. Analysis of cell wall chitin was assessed using calcofluor white staining, as described previously (21). Briefly, cells were fixed in 3.7% formaldehyde, standardized to 1 x 10^6^ cells/ml in PBS, and stained for 5 minutes at 25°C with 1μg/ml of calcofluor white (Sigma Aldrich, St. Louis, MO). Stained cells were pelleted and resuspended in PBS. 10,000 cells for each sample were analyzed using a BD LSR II H4710 or H1160 (Becton, Dickinson, Hercules, CA), and data were analyzed using Flowjo v10 software (Treestar, Ashland, OR).

Analysis of cell morphology based on the ploidy of the cell was performed using a BD FACSAria II cell sorter. *C. neoformans* cells were isolated from the lungs of mice 14 days after infection as described above. The cells were then passed over a 20 µm filter to remove the titan cells from the sample, with the flow-through typically containing a purified population that is >98% typical sized cells (73). Log phase cells were grown by inoculating YPD broth medium and incubating the culture overnight at 250 rpm. Stationary phase cells were grown similarly but the cultures were incubated for 96 hours. All samples were fixed and stained with PI as described above. Cells were gated for singlets and sorted based on PI fluorescence with at least 400,000 cells sorted into each population. A minimum of 100 cells in each of the resulting populations were analyzed for budding morphology using a Zeiss AxioImager microscope (Carl Zeiss, Inc., Thornwood, NY) equipped with ZenBlue2 software (Carl Zeiss, Inc., Thornwood, NY).

### Fluorescence microscopy

Prior to live cell fluorescence imaging, overnight cultures were pelleted at 15,000 rpm and washed with complete minimal media (CMM) and resuspended in CMM. For analysis of log phase and stationary phase typical sized cells, cells were spotted and imaged on a 2% agarose patch supplemented with CMM and imaged on a Zeiss AxioImager (Carl Zeiss, Inc., Thornwood, NY). For analysis of *in vivo* titan cells, cells were resuspended in CMM cells and 500 µl were placed in a 35mm glass bottom microwell petri dish 14mm microwell dish (MatTek Corporation, Ashland, MA). *In vivo* titans were then imaged on a Zeiss LSM 800 confocal laser scanning microscope (Carl Zeiss, Inc., Thornwood, NY) or a Zeiss Cell Observer SD spinning disk microscope (Carl Zeiss, Inc., Thornwood, NY). Images were analyzed using ZenBlue2 software (Carl Zeiss, Inc., Thornwood, NY).

### Stationary phase analysis

To analyze the ability of cells to enter stationary phase overnight cultures were diluted to a final concentration of 1x10^5^ cells/ml in YPD and incubated at 30°C. At 1, 2, 3, 4, and 5 days after starting the culture, 1 ml of cell suspension was harvested. Cell concentration was enumerated using a hemocytometer and cell morphology was determined by analysis of at least 100 cells on a Leica DM750 microscope (Leica Biosystems, Wetzlar, Germany). For ploidy analysis, cells were fixed with 70% ethanol, stained with PI, and analyzed by flow cytometry as described above. To assess cells after stationary phase release, cells were grown for 96 hours under nutrient starvation. Cells were then centrifuged to pellet the cells, washed with PBS twice, and resuspended to a final concentration of 2.5x10^5^ cells/ml in YPD broth and incubated at 30°C with shaking at 250 rpm. The cell morphology was determined by analysis of at least 100 cells per time point on a Leica DM650 microscope (Leica Biosystems, Wetzlar, Germany).

### Spot Assays

Cultures were washed two times with PBS, resuspended in PBS, and diluted to 2x10^6^ cells/ml. The resulting cells were serially diluted ten-fold and 5 µl of each dilution was spotted onto YPD plates. Plates were incubated at 30°C and growth was assessed after 48 hours of incubation.

### RNA purification and RT-qPCR

For RT-qPCR analysis of synchronized cell populations, stationary phase cells were generated after 72 hours incubation as described above. For comparison of *CLN1* RNA level in titan and typical sized cells, cell concentration was normalized prior to initiation of the RNA extraction protocol. RNA extraction was performed as previously described (74). Cells were pelleted, frozen in liquid nitrogen, and stored at -80 °C. The frozen pellet was lyophilized and vortexed to powder. RNA was extracted using a modified PureZOL RNA isolation reagent/Qiagen procedure. Cells were lysed in 700 µl PureZOL RNA isolation reagent (Bio-Rad, Hercules, CA) and incubated for 5 minutes at room temperature. Then 140 µl chloroform was added, mixed thoroughly, and centrifuged at 20,000 x g for 5 minutes at room temperature. The aqueous phase was separated and mixed with an equal volume of 70% ethanol and immediately applied to an RNeasy Mini Spin Column (Qiagen, Valencia, CA). RNA was isolated according to the manufacturer’s protocol. The quality and quantity of RNA were measured using a NanoDrop spectrometer (Thermo Scientific). The iScript one-step RT-PCR Kit with SYBR Green (Bio-Rad, Hercules, CA) was used for RT-qPCR following the manufacturer’s protocol. RT-qPCR was performed in an iQ5 real-time PCR detection system (Bio-Rad, Hercules, CA). Gene expression levels were normalized using the endogenous control genes *TEF1* or *ACT1*. Relative copy levels were determined using the comparative threshold cycle (*C*_T_) method (75).

### Co-IP and near-infrared (NIR) western blot analysis

The *CLN1-His*_6_ *CDK1-Myc* strain was cultured in 50 ml YPD liquid medium for 1.5 days at 30°C (250 rpm) to reach stationary phase. The culture was diluted 1:70 with fresh YPD and 0 min, 15 min, 30 min, 45 min, 60 min, 75 min, or 90 min time points were collected as described above. Cells were pelleted, frozen in liquid nitrogen, and stored at -80°C. The frozen pellet was lyophilized, vortexed to powder, and suspended in lysis buffer. Cdk1 and its associated proteins were co-purified using the Pierce™ c-Myc-Tag IP/Co-IP Kit (Thermo Scientific, Rockford, IL). Cell lysates from the *CDK1-Myc* strain were used as a negative control. Cln1-His_6_ and its associated proteins were co-purified following the Ni-NTA Spin Kit procedure (Qiagen, Valencia, CA). Cell lysate from the *CLN1-His*_6_ strain was used as a negative control. The purified proteins were separated by SDS/4–15% Mini-PROTEAN® TGX™ Precast Protein Gels (Bio-Rad, Hercules, CA), then transferred to an Immobilon-FL PVDF membrane (Merck Millipore Ltd., Tullagreen, Ireland) using a Trans-Blot® Turbo™ Transfer System (Bio-Rad, Hercules, CA) for western blot analysis. The blots were incubated overnight at 4°C with the primary antibodies: His-Tag (D3I1O) XP® Rabbit mAb (1/1,000 dilution) (Cell Signaling Technology, Danvers, MA) and Myc-Tag (9B11) Mouse mAb (1/1,000 dilution) (Cell Signaling Technology, Danvers, MA), then incubated 1 h at room temperature with the secondary antibodies: IRDye® 800CW Goat anti-Rabbit IgG (H + L) (1/20,000 dilution) (LI-COR, Lincoln, NE) and IRDye® 680CW Goat anti-Mouse IgG (H + L) (1/20,000 dilution) (LI-COR, Lincoln, NE). Detection was performed using an Odyssey Fc infrared imaging system (LI-COR, Lincoln, NE).

### Kinase activity assay

The kinase activity assay was performed as previously described with the following modifications (76). The *CLN1-His*_6_ *CDK1-Myc* strain was cultured in 50 ml YPD liquid medium for 1.5 days at 30°C, 250 rpm to reach stationary phase. The culture was then diluted 1:70 with fresh YPD and cultured an additional 15 min or 45 min. Cells were pelleted, frozen in liquid nitrogen, and stored at -80°C, lyophilized, vortexed to powder, suspended in lysis buffer, and 2 mg of total protein was incubated with 15 µl Goat anti-Myc Antibody Tag Agarose Immobilized (Bethyl, Montgomery, TX) at 4°C overnight. The beads were washed three times with cold lysis buffer on Pierce™ Spin Columns - Screw Cap (Thermo Scientific, Rockford, IL) and then suspended in 15 µl 1x kinase buffer. The kinase activity assay was performed with 4 µl beads in a 384-well plate using the ADP-Glo™ Kinase Assay (Promega, Madison, WI). The luminescence of each well was read using a Synergy™ H1 Hybrid Multi-Mode Microplate Reader (BioTek Instruments, Winooski, VT). Data was collected using Gen5™ Data Analysis Software (BioTek Instruments, Winooski, VT) and the 1x kinase buffer was used as a negative control.

### Phylogenetic and motif analysis of cyclins

Multiple sequence alignment of the cyclins was performed using ClustalX2 (77). The neighbor-joining tree was created in MEGA6.06 using the Jones-Taylor-Thornton (JTT) model. Conserved motifs of cyclins were analyzed using MEME Suite 4.10.1 (http://meme-suite.org/index.html). The MEME program was employed using the following parameters: Zero or one occurrence per sequence, 10 motifs, motif width set between 6 and 50, site number of each motif set between 2 and 600.

### Complementation of *S. cerevisiae* cyclin mutants

An overlap PCR product was created with the *S. cerevisiae GPD1* promoter (865 bp), cDNA of *C. neoformans CLN1* generated by reverse transcription, and the *S. cerevisiae ACT* terminator (319 bp). The PCR product was digested with *Hin*dIII and *Xba*I and then inserted into pRS316 (*URA3* marker) or pRS428 (*ADE1* marker) to generate pRS316-*P_GPD1_*-*CLN1* or pRS428-*P_GPD1_*-*CLN1.* pRS316-*P_GPD1_*-*CLN1* or pRS428-*P_GPD1_*-*CLN1* was introduced into *S. cerevisiae cln1 cln2 cln3* triple mutant CWY364 (*MAT**a** cln1△cln2△cln3△leu::GAL-CLN3::ade1 his2 trp1-1 ura3 △ns bar1△*), *S. cerevisiae clb3△clb4△clb5△clb6△* quadruple mutant K3418F (*MAT**a** ade2-1can1-100 his3-11,15 leu2 trp1-1 ura3-1 clb3::TRP1 clb4::HIS3 clb5::hisG clb6::LEU2 TRP1::GAL-CLB5*), or *S. cerevisiae clb1 clb2 clb3 clb4* quadruple mutant YS108 (*MATα GAL1::CLB1 (LEU2) clb1::URA3 clb2::LEU2 clb3::TRP1 clb4::HIS2 ade1*) by the LiAc/SS Carrier DNA/PEG method, respectively (78).

Transformants were selected on uracil-free or adenine-free media with galactose, and then tested for growth on YPD medium without galactose.

### *In vitro* titan cell formation

Titan cells were generated *in vitro* as previously described (43). Briefly, cells were grown in 10 ml of YPD for 22 hours at 30°C. Cells were then washed with minimal media, resuspended at a final concentration of 1x10^6^ cells/ml in minimal media in a 1.5 ml tube and grown at 30°C with 800 rpm in an Eppendorf Thermomixer (Eppendorf, Hamburg, Germany). Cell size was analyzed after 4 days. For analysis of *CLN1* expression during *in vitro* titan cell formation, 25 μM copper (Cu^2+^) or 400 μM of the copper chelator bathocuproinedisulfonic acid (BCS) was added during overnight growth and/or during incubation in minimal media.

### Statistical analysis

Microsoft Excel and GraphPad Prism version 9 were used for statistical analysis. Data sets were analyzed for normality and global tests were performed by ANOVA or Kruskal Wallis with appropriate corrections for multiple comparisons. Pairwise comparisons were carried out by Student’s t-test with Welch’s correction for multiple comparisons. P-values ≤ 0.05 were considered statistically significant.

## Acknowledgements

We thank Dr. Curt Wittenberg (the Scripps Research Institute), Dr. David Stuart (University of Alberta), and Dr. Steven Reed (the Scripps Research Institute) for providing *S. cerevisiae* strains and Dr. Joseph Heitman and Dr. Lukasz Kozubowski for providing *C. neoformans* strains. We would also like to thank Dr. Emma Robertson for technical support, and the University of Minnesota Flow Cytometry Core Facility and the University of Minnesota Imaging Center for instrumentation. This work was supported by National Institutes of Health (NIH) grants R01AI080275 and R01AI134636 to KN, SA was supported by an NIH IRADCA fellowship via NIH grant K12GM119955 and MD was supported by an NIH MSTP fellowship via NIH grant T32GM008244.

## Author Contributions

Conceived and designed the experiments: SA, ZL, MSF and KN. Performed the experiments: SA, ZL, MSF, and JMY. Analyzed the data: SA, ZL, MSF, MD, VT, JMY, SRF and KN. Contributed reagents/materials/analysis tools: KN. Wrote the paper: SA, ZL, and KN.

